# Vancomycin tolerance and dispersion of dual species biofilms of *Clostridioides difficile* and Vancomycin-resistant *Enterococcus faecium*

**DOI:** 10.64898/2026.03.09.710618

**Authors:** Holly R. Neubauer, Sarah Joseph, Ishtiaq Ahmad, Peter T. McKenney

## Abstract

**Objectives:** Biofilms are the dominant mode of bacterial life. The gut microbiota itself has characteristics of a biofilm that grows on the intestinal mucosa. *C. difficile* and VRE are commonly co-isolated from patients but biofilm formation has not been studied in a multi-species context. Here we study the interactions between *C. difficile* and VRE in surface adherent community.

**Results:** We found that VRE inhibits *C. difficile* biofilm formation in dual-species culture in the presence of excess glucose. Robust dual-species biofilms were produced when the carbon source was changed to a non-fermentable sugar such as fucose and xylose. We observed a high level of vancomycin tolerance in *C. difficile* biofilms that was not affected by the presence of VRE. Finally we also found that a nutrient step-change is sufficient to induce dispersion of single and dual-species biofilms.

**Conclusions:** VRE can inhibit the development of *C. difficile* biofilms in the presence of a fermentable carbon source. VRE does not appear to affect vancomycin tolerance or nutrient-induced dispersion of *C. difficile* biofilms.

**Highlights:**

- VRE inhibits *C. difficile* biofilm formation in the presence of fermentable glucose.
- Stable VRE – *C. difficile* biofilms are formed by managing the available carbon source.
- VRE does not affect *C. difficile* vancomycin tolerance in this model.
- A 10-fold increase in available nutrients is sufficient to induce biofilm dispersion in *C. difficile* and VRE.

## Introduction

*Clostridiodes difficile* is a leading cause of nosocomial infection that is commonly co-isolated with Vancomycin-resistant *Enterococcus faecium* (VRE)^1^. Enterococci generally enhance *C. difficile* virulence^2,3^ and also predispose cancer patients to *C. difficile*^4^. In the case of *C. difficile* the production of dormant stress resistant spores complicates treatment and, in part, leads to high levels of transmission and recurrence^5,6^. Both *C. difficile* and VRE are also capable of forming biofilms^7,8^. Biofilms are adherent communities of bacteria that form on surfaces and have been proposed as a mechanism that may contribute to recurrent *C. difficile* infection^9,10^.

Biofilms are a ubiquitous growth mode of bacteria^11^. Cells adhere to each other and to a surface, forming small microcolonies which then expand in place and produce an extracellular polymer matrix^12,13^. The biofilm surface can form a protective barrier between interior cells and the outside environment^14^. Biofilm cells are protected from harsh conditions such as immune recognition, antibiotic treatment and competition with other bacteria^15^. In nature bacteria create multispecies biofilms, in which bacteria can exchange nutrients, exchange genes, and harvest host-associated metabolites^16^. The gut microbiota in healthy humans and animals has characteristics of a biofilm^17^, which are disrupted in models of infectious disease^18^. This suggests that understanding interactions between *C. difficile* and VRE in the biofilm mode of growth represents an important knowledge gap.

One difficulty in multi species biofilm models is developing a compatible growth medium that supports multiple species. Our previous work found that VRE metabolizes modest amounts of glucose, fructose or trehalose into acid sufficient to inhibit *C. difficile*^19^. In this work we established a long-term dual-species biofilm model of *C. difficile* and VRE. We used this assay to characterize vancomycin tolerance in mono and dual-species biofilms. VRE did not appear to affect vancomycin tolerance of *C. difficile* under these conditions. Finally, we characterized nutrient step-change induced biofilm dispersion in mono and dual-species biofilms. Biofilm dispersion in both species appeared to be regulated independently with no obvious inter-species effect in dual species biofilms.

## Materials and Methods

### Strains and Growth Conditions

All experiments were conducted in an anaerobic chamber (Coy Laboratory Products), with an atmosphere of 90% N_2_, 5% CO_2_ and 5% H_2_. *C. difficile* (VPI 10463) and Vancomycin-resistant *Enterococcus faecium* (VRE)(ATCC: 700223) were grown in BHI liquid medium supplemented with 0.5% yeast and 0.1% cysteine overnight to an OD600 of 0.3-0.8. Cultures were OD600 normalized and diluted 1:10 into sporulation media (SM). Dual species cultures were inoculated at a ratio of 1 VRE to 10 *C. difficile* to compensate for the faster growth rate of VRE. All experiments and were plated on *C. difficile* selective agar (BHIS + 0.1% taurocholate, 250 µg/mL D-cycloserine, 16 ug/mL cefoxitin) and/or VRE selective agar (Enterococcosel + 8 µg/mL vancomycin). Inoculums were cross plated to ensure there was no cross contamination between *C. difficile* and VRE. For biofilms, 24 well, non-coated polystyrene plates were used for all experiments. The innermost 8 wells of the 24 well plates were used to grow biofilms and the remaining periphery wells were filled ½-⅔ full with sterile water to prevent dehydration and edge effects. Biofilms were incubated anaerobically at 37C. For biofilms with fibrinogen coating, 100 µL of 5mg/mL human fibrinogen was added to each well and left overnight or until dry, before reducing in the anaerobic chamber.

### Confocal Imaging

Biofilms were stained with BacLight (Invitrogen) then fixed with 4% PFA in their 24 well plates for microscopy. A Leica-Sp5 confocal scanning microscope with long working distance 10x objective was used for imaging. Wavelengths of 490 / 480 excitation and emission respectively, with 30% laser power. 10x zoom was used to visualize individual biofilm fields and 3-D acquired using 3.00 micron z-steps. LAS X application version 3.7.4.23463 was used to visualize the z-stacks and generate images in Figure 1. Comstat 2.1 analysis was performed using default settings to analyze biomass, average thickness and surface area to biovolume. Three fields were collected from each well and averaged for each data point.

**Figure 1:**
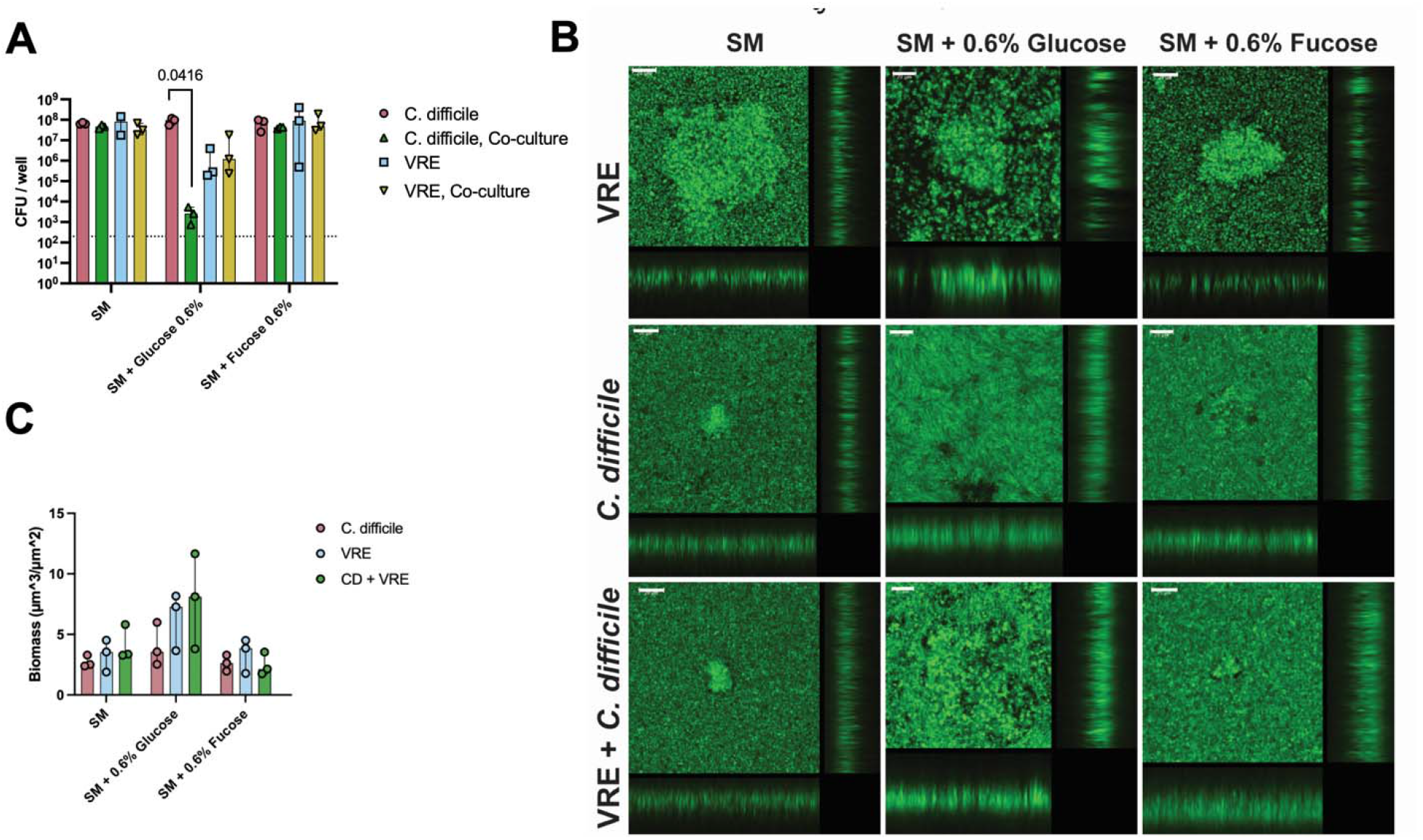
VRE inhibits *C. difficile* in biofilm cultures with added glucose. A) CFU counts of biofilms grown for 48 hours in single or dual species biofilms grown in the indicated medium. Plates were pre-coated with fibrinogen to aid in adhesion. n = 3 biological replicates. Statistics = t-test with Welch’s correction single vs. dual species culture. B) Laser scanning confocal microscopy of with z-projection of 48 hour biofilms. C) Biomass of biofilms quantified using COMSTAT.

### Time course biofilms

Biofilms were grown up to 5 days for each replicate experiment and were harvested on days 2, 3, and 5. One set of replicates remained in a static culture that was not disturbed, the other set was in a static culture with a media replacement every 24 hours. Biofilms were rinsed once with PBS and resuspended in 1.0 mL of PBS. Wells were scraped in all directions and pipetted up in down multiple times to ensure attached cells were resuspended followed by vigorous vortexing to dissociate individual cells. Single cell suspension was verified by light microscopy. Samples were serial diluted and plated on *C. difficile* and VRE selective plates.

### Vancomycin Biofilm Tolerance

Biofilms were grown for 4 days in SM + 125 µM DCA without medium changes. On the fourth day, medium was replaced with fresh SM + 125 µM DCA + 0, 32, or 64 µg/mL vancomycin. Biofilms were incubated overnight and processed as described above for CFU’s.

### Nutrient Step Dispersion

Biofilms were grown in 10% SM + 125 µM DCA for 5 days without medium changes. On day 5 biofilms were washed for 1 hour with fresh 10% SM + 125 µM DCA followed by replacement with 100% SM for 1 hour. Supernatant and biofilms were harvested as described above.

### Statistical analysis

All graphs and statistical analysis was performed in GraphPad Prism 10. All statistical tests are stated within figure legends.

## Results

### *C. difficile* and VRE can form stable biofilms and macrocolonies within 48 hours

Our primary goal was to establish a dual-species biofilm model of growth between *C. difficile* and VRE. We began this optimization by growing both species 1:1 in non-treated polystyrene plates in sporulation media (SM), a medium with no added carbon source^20^. We added 125 µM added deoxycholic acid (DCA) to the SM to stimulate biofilm adherence in both species^8,21^ and pre-coated the plates with human fibrinogen^22^. In Gram positive bacteria, excess glucose is generally added to media to induce biofilm formation^7^. VRE converts glucose to acid and drops the pH of liquid media to a point that is toxic to *C. difficile*^19^. To test the effect of carbon source on biofilm formation we added glucose or fucose, which is not acidified by VRE^19,23^, to the media. We found that VRE inhibits *C. difficile* growth in the presence of glucose, whereas in SM + xylose or SM + fucose growth of *C. difficile* and VRE are supported (Figure 1A). We performed confocal imaging on 48 hour biofilms with SYTO-9 stain and found the formation of macrocolonies in mono- and dual-species biofilms in each carbon source (Figure 1B). Quantitative measurement of biofilms using COMSTAT^24^, did not reveal significant changes in biofilm thickness or surface area (Figure 1C). These data suggest *C. difficile* and VRE can form a stable dual-species biofilms in the presence of xylose and fucose and in unsupplemented SM + DCA.

### Long-term *C. difficile* - VRE biofilms

We expanded our initial 48 hour biofilm model to 5 days to analyze longer term interactions of *C. difficile* and VRE. Others have demonstrated that media changes performed every 24 hours may be beneficial to promote biofilm growth and reduce build up of waste products^25^. We analyzed biofilm CFUs on days 2, 3 and 5 following inoculation. In ‘batch’ culture of unsupplemented SM + DCA without media changes, a stable and durable dual-species biofilm was formed with a not significant trend towards lower *C. difficile* CFUs in co-culture (Figure 2A). Addition of 0.6% glucose to SM resulted in a statistically significant reduction in *C. difficile* CFUs at days 2, 3 and 5 in co-culture. VRE levels were not affected by co-culture with *C. difficile*, however, total VRE CFUs were about 1-log lower in the presence of added glucose (Figure 2A). The reduction in *C. difficile* and VRE CFU levels in SM + Glucose were likely the result of acid production by VRE. In SM + glucose cultures containing VRE, the pH was generally ∼ 6.0 (Figure 2B), which is growth inhibitory towards *C. difficile*^19,26^ and limiting towards VRE^19^. Changing the medium every 24 hours resulted in a recovery of *C. difficile* CFUs by Day 5 when in co-culture with VRE (Figure 2C), without a concomitant rise in pH (Figure 2D). In these conditions, *C. difficile* and VRE encounter fresh medium at a neutral pH daily. This should allow for repeated daily growth of *C. difficile*. The growth medium for these experiments contained 125 μM of DCA, which stimulates biofilm formation in both *C. difficile*^21^ and VRE^8^. Deoxycholic acid also stimulates spore germination^27^. Thus, it is likely that the recovery of *C. difficile* in this long-term co-culture biofilm with daily medium change is driven by proliferation of dormant vegetative cells and germination of spores. It is also possible that in these conditions *C. difficile* may adapt to repeated exposure to competition with VRE and repeated acidification of the local environment.

**Figure 2:**
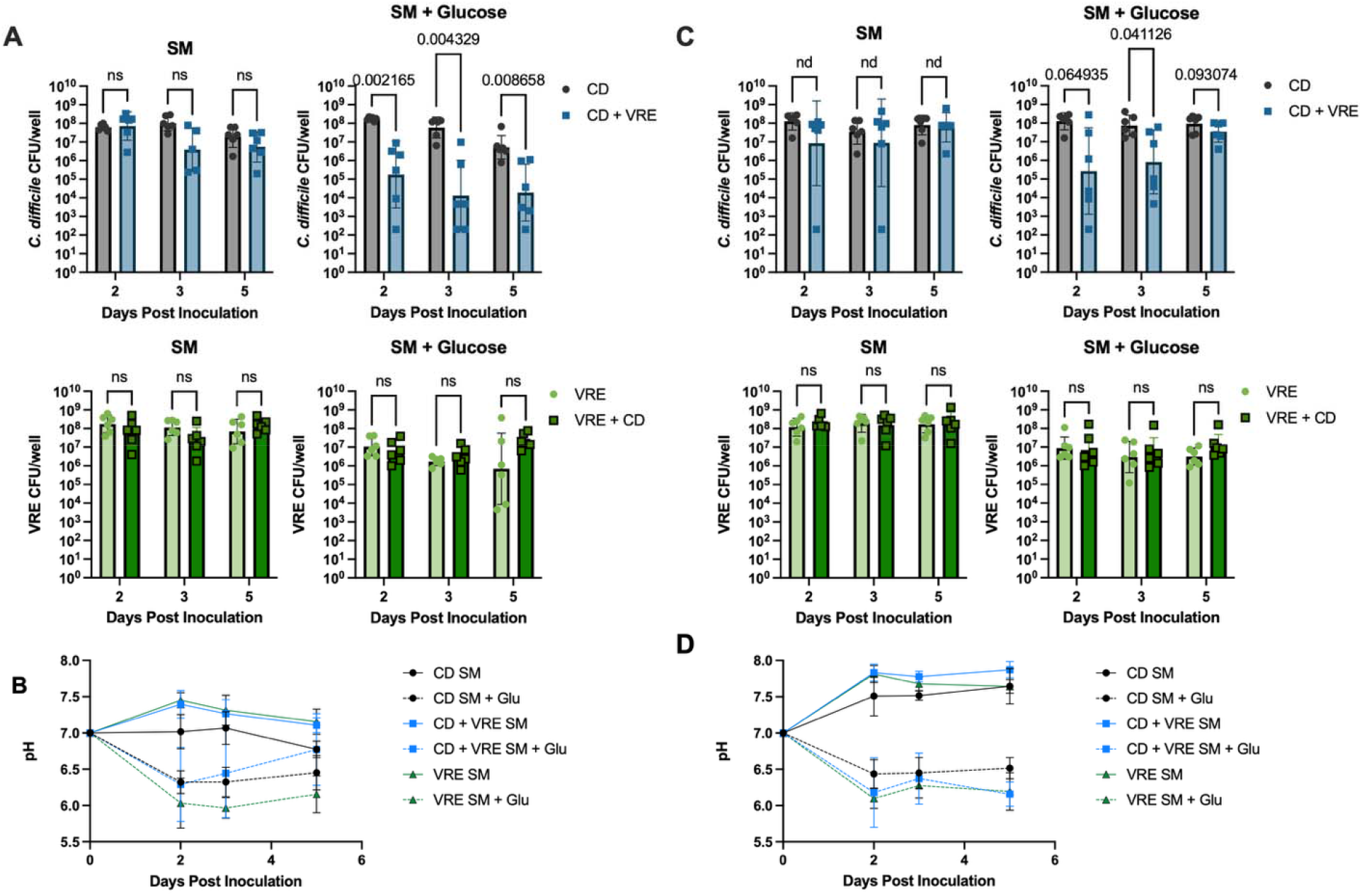
Medium change allows *C. difficile* to grow with VRE in the presence of excess glucose. **A)** CFU counts of single and dual species biofilms grown in SM or SM + 0.6% glucose in batch format without medium change. **B)** pH of medium from A following overnight growth.**C)** CFU counts of single and dual species biofilms grown in SM or SM + 0.6% glucose with daily medium change. **D)** pH of medium from C following overnight growth. Statistics: t-test with Welch’s correction. n = 6 biological replicates.

### *C. difficile* biofilms have a significant vancomycin tolerance

*Enterococcus faecalis* OG1RF, which is vancomycin sensitive, protected *C. difficile* from vancomycin treatment during biofilm growth^2^. We tested if VRE, which is resistant to vancomycin, can also have a protective effect on *C. difficile*. Previous studies suggested that *C. difficile* tolerance to vancomycin is enhanced in biofilms^7,21^. We first performed a dose-finding experiment of late log phase vegetative *C. difficile*. We observed a 1 log reduction at 16 µg/mL vancomycin with overnight exposure (Figure 4A). We then went on to treat 4-day biofilms with 32 and 64 µg/mL vancomycin overnight (Figure 4B). We observed a significant decrease in *C. difficile* CFUs at 64 µg/mL vancomycin in single and dual species biofilms and a slight, but significant, decrease in *C. difficile* single species biofilms at 32 µg/mL. We did not observe a significant difference when comparing *C. difficile* survival in single and dual species biofilms.

### Nutrient step-change induces biofilm dispersion

The final step of the biofilm development life cycle is dispersion, a programmed disassembly that allows cells to move to a new location. Dispersion has not been characterized in *C. difficile* or VRE. In *Pseudomonas aeruginosa*, dispersion of mature biofilms can be induced by a sudden step-change in available nutrients^25^. We grew biofilms for 4 days in 10% SM + DCA with medium change every 24 hours. On day 5 we replaced the media with 100% SM without DCA and plated the released cells in the supernatant and the attached biofilm samples after 60 minutes. We observed a consistent increase in the raw CFU content of the supernatant which was significant for *C. difficile* grown alone (Figure 4A) and in dual species biofilms (Figure 4C) and for significant for VRE in dual species biofilms (Figure 4C). When we normalized the CFU counts as the percentage of released cells in total, the percentage of released cells in 100% SM was significantly increased when compared to 10% SM for both species in mono and dual-species biofilms (Figure 4D). These data suggest that nutrient step-change is sufficient to induce biofilm dispersion in both species. We did not observe a significant difference for *C. difficile* or VRE when comparing the percent released in 100% SM between mono species or dual-species biofilms, indicating that VRE and *C. difficile* do not influence one another during dispersion under these conditions. We note that the VRE strain used in this study, *E. faecium* ATCC 700221, forms weakly adherent biofilms. We hypothesize that strains of enterococci that form strongly adherent biofilms, such as *E. faecalis* OG1RF, may affect the dispersion of *C. difficile*.

## Discussion

Taken together we have shown that *C. difficile* and VRE are capable of forming dual species biofilms, provided that the medium does not contain an excess of sugar that is fermentable by VRE (Figure 1). In our dual-species biofilms we did not observe a significant difference between SM and SM supplemented with a non-fermentable carbon source such as fucose or xylose (Figure 1). In long term dual species batch culture without medium change, VRE acidified media containing glucose and *C. difficile* did not proliferate (Figure 2A & 2B). Daily medium change allowed *C. difficile* to overcome the acidification mediated inhibition of VRE in a glucose rich medium (Figure 2C & 2D). With medium change, the final pH of the bulk liquid in dual species culture with added glucose after overnight growth was between 6.2 and 6.4 (Figure 2D) on days 2, 3 and 5. In spite of this growth inhibitory pH, *C. difficile* was able to reach CFU levels comparable to single species culture. It is likely that *C. difficile* undergoes daily cycles of proliferation when fresh pH neutral medium is added each day. Repeated daily cycles of growth and stationary phase may more accurately reflect the natural ecology of *C. difficile* which must tune its lifecycle to the circadian rhythms of the host.

We also described high levels of vancomycin tolerance for both *C. difficile* VPI 10463 vegetative cells grown in SM medium and established biofilms (Figure 3). These data suggest that *C. difficile* biofilms have significantly increased tolerance to vancomycin, as has been observed previously^7,21^. We did not observe protection of *C. difficile* by VRE in the presence of vancomycin (Figure 3B). We note that a previous study used a different strain of *C. difficile* (CD196) and vancomycin sensitive *E. faecalis* OG1RF in a different liquid medium (BHI) to characterize protection of *C. difficile* in co-culture^2^. Vancomycin binds to terminal D-Ala-D-Ala motifs on lipid II^28^. It is possible that vancomycin sensitive *E. faecalis* OG1RF, with its abundance of free D-Ala-D-Ala lipid II, acted partly as a ‘false target’ for vancomycin^29^ in that experiment. Binding to *E. faecalis* stem peptides could potentially lower the effective concentration of free vancomycin available to bind and kill *C. difficile* in co-culture. In our co-culture biofilms here the lipid II of VRE (*E. faecium* ATCC 700221), are converted to D-Ala-D-Lac in the presence of vancomycin which prevents binding of the antibiotic^28^ and may negate any protective effects of co-culture. In the future it may be useful to directly compare the effects of co-culture with congenic vancomycin sensitive and vancomycin resistant *Enterococcus* strains to determine if the false target model applies to multi-species populations with *C. difficile*. Vancomycin-intermedia *Staphylococcus aureus* (VISA) strains produce an excess of weakly cross-linked peptidoglycan and a thicker cell wall which has been proposed to act as a ‘net’ of free D-Ala-D-Ala stem peptides to prevent vancomycin from reaching lipid II at the membrane^29^. Inter-species interactions that affect production of lipid II or cross-linking of mature peptidoglycan have the potential to impact glycopeptide susceptibility in mixed populations.

**Figure 3:**
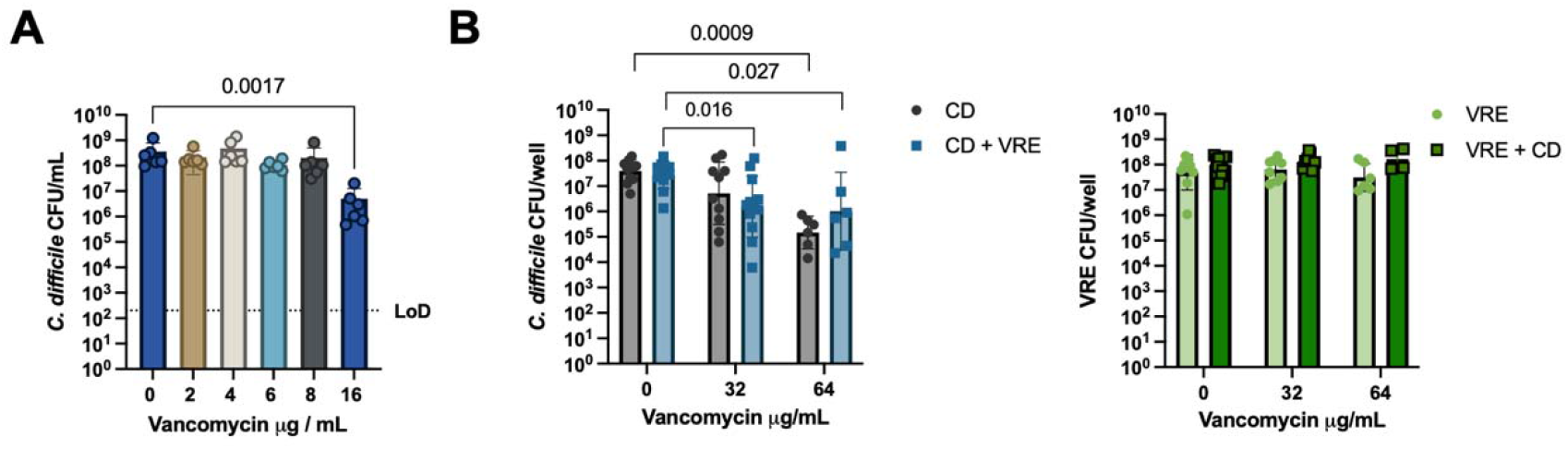
*C. difficile* biofilm vancomycin tolerance is not affected by VRE. A) CFUs of *C. diffcile* late mid-log culture exposed to vancomycin for 24 hours. n = 6 biological replicates.Statistics: Kruskal-Wallace One-way ANOVA versus untreated. B) Single and dual species biofilm CFUs of 48 hour biofilms exposed to the indicated concentration of vancomycin for 24 hours. n = 6-12 biological replicates per condition. Statistics: Kruskal-Wallace One-way ANOVA versus untreated.

Finally, a sudden 10-fold increase in nutrient availability appears to be sufficient to trigger biofilm dispersion in mono- and dual-species biofilms of *C. difficile* and VRE (Figure 4). It will be interesting to determine the effects of metabolites from the intestinal nice such as bile acids and short chain fatty acids on *C. difficile* dispersion. Given that secondary bile acids promote biofilm formation^8,21^, it is likely that they would inhibit dispersion. Secondary bile acid levels fall in the colon after antibiotic treatment^30^. Dispersion of *C. difficile* following antibiotic treatment in colonic biofilms, which would release motile vegetative toxin producing cells, may play an important role in pathogenesis. In other systems, induction of dispersion is sufficient to eliminate the elevated antibiotics tolerance of biofilms^31^. Induction of dispersion combined with antibiotic treatment could increase the efficacy of antibacterial drugs against biofilms. This approach would come with the risk of inducing systemic infection^32^. If the regulatory pathways of biofilm dispersion are characterized in detail, it may be possible to genetically uncouple the desired effect of lowering antibiotic tolerance from the potentially dangerous disassembly of the biofilm.

**Figure 4:**
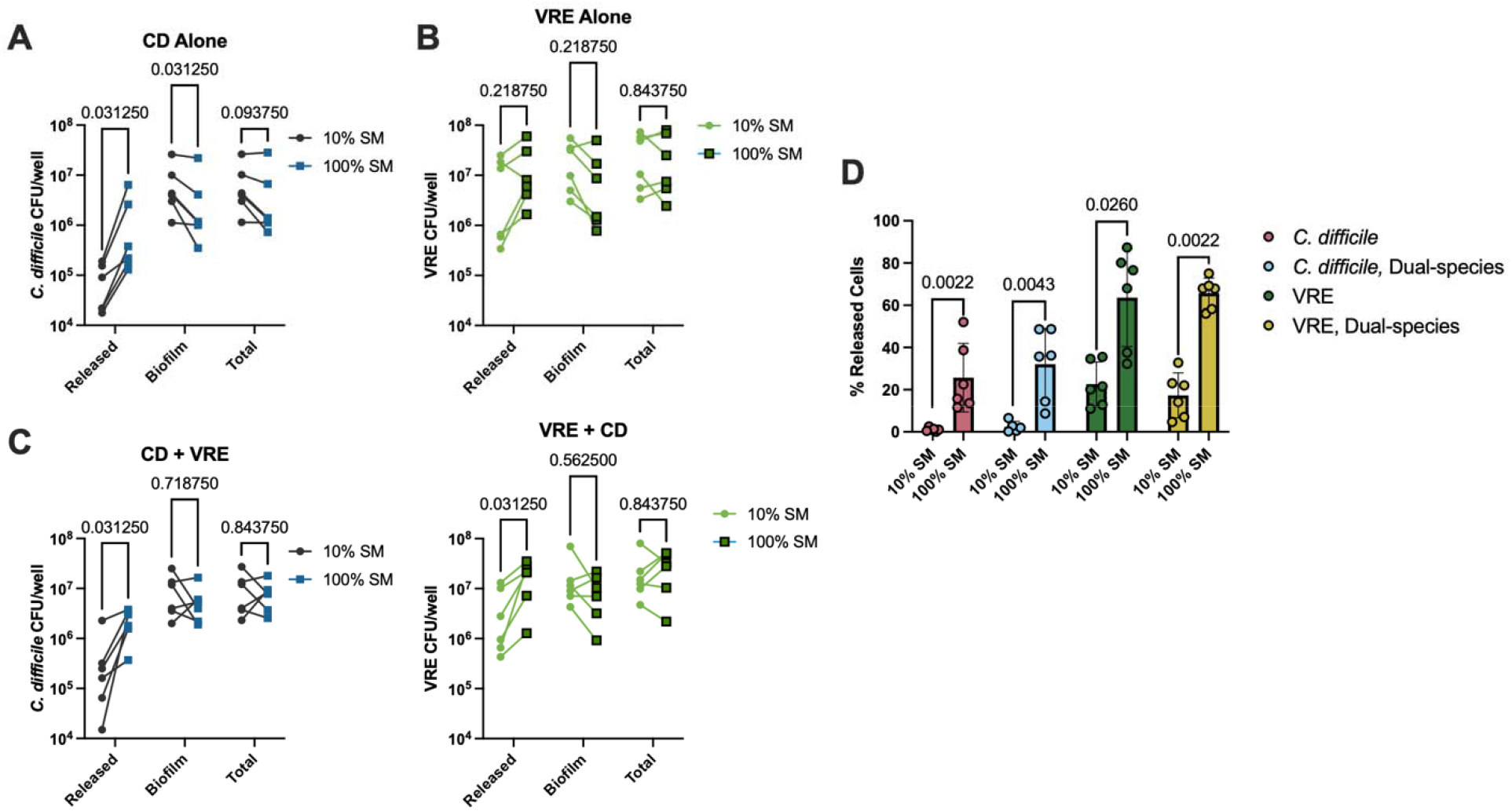
VRE does not affect nutrient step-change induced dispersion of *C. difficile*. 48 hour biofilms grown in 10% SM were exposed to 100% SM for 1 hour then enumerated by selective plating. Released cells represent the cells recovered in the liquid medium after 1 hour. Biofilm cells represent cells recovered attached to the plate. **A)** *C. difficile* single species biofilm. **B)** VRE single species biofilm. **C)** *C. difficile* and VRE dual species biofilm. **D)** Data from A, B and C normalized as percent released cells. Statistics: t-test with Welch’s correction. n = 5-6 biological replicates.

The ability to selectively switch off elevated biofilm antibiotic tolerance could increase the efficacy of existing antibacterial drugs.

## Acknowledgements

We thank Victoria Oladosu for help with confocal microscopy, David Davies and Claudia Marques for help with adaptation of the nutrient step-change assay and members of the Binghamton Biofilm Research Center for input on experimental design and troubleshooting. This work was supported by startup funds from Binghamton University a grant from the Analytical & Diagnostics Laboratory at Binghamton University and NIH R21AI171634.

## References

1. Tickler, I. A. et al. Presence of Clostridioides difficile and multidrug-resistant healthcare-associated pathogens in stool specimens from hospitalized patients in the USA. J. Hosp. Infect. 106, 179–185 (2020).

2. Smith, A. B. et al. Enterococci enhance Clostridioides difficile pathogenesis. Nature 611, 780–786 (2022).

3. Keith, J. et al. Impact of antibiotic-resistant bacteria in the mouse intestine on immune activation and Clostridioides difficile infection. Infect. Immun. (2020) doi:10.1128/IAI.00362-19.

4. Lee, Y. J. et al. Protective Factors in the Intestinal Microbiome Against Clostridium difficile Infection in Recipients of Allogeneic Hematopoietic Stem Cell Transplantation. J. Infect. Dis. 215, 1117–1123 (2017).

5. Shen, A. Clostridioides difficile Spore Formation and Germination: New Insights and Opportunities for Intervention. Annu. Rev. Microbiol. 74, 545–566 (2020).

6. Castro-Córdova, P. et al. Entry of spores into intestinal epithelial cells contributes to recurrence of Clostridioides difficile infection. Nat. Commun. 12, 1140 (2021).

7. Đapa, T. et al. Multiple factors modulate biofilm formation by the anaerobic pathogen Clostridium difficile. J. Bacteriol. 195, 545–555 (2013).

8. McKenney, P. T. et al. Intestinal Bile Acids Induce a Morphotype Switch in Vancomycin-Resistant Enterococcus that Facilitates Intestinal Colonization. Cell Host Microbe 25, 695-705.e5 (2019).

9. Normington, C. et al. Biofilms harbour Clostridioides difficile, serving as a reservoir for recurrent infection. NPJ Biofilms Microbiomes 7, 16 (2021).

10. Meza-Torres, J., Auria, E., Dupuy, B. & Tremblay, Y. D. N. Wolf in Sheep’s Clothing: Clostridioides difficile Biofilm as a Reservoir for Recurrent Infections. Microorganisms 9, (2021).

11. Costerton, W. BACTERIAL BIOFILMS IN NATlJRE AND DISEASE. Ann. Rev. Microbiol. 41, 435–464 (1987).

12. Tremblay, Y. D. & Dupuy, B. The blueprint for building a biofilm the Clostridioides difficile way. Curr. Opin. Microbiol. 66, 39–45 (2022).

13. Ronish, L. A., Biswas, B., Bauer, R. M., Jacob, M. E. & Piepenbrink, K. H. The role of extracellular structures in Clostridioides difficile biofilm formation. Anaerobe 88, 102873 (2024).

14. Pantaléon, V., Bouttier, S., Soavelomandroso, A. P., Janoir, C. & Candela, T. Biofilms of Clostridium species. Anaerobe 30, 193–198 (2014).

15. Arciola, C. R., Campoccia, D. & Montanaro, L. Implant infections: adhesion, biofilm formation and immune evasion. Nat. Rev. Microbiol. 16, 397–409 (2018).

16. Stoodley, P., Sauer, K., Davies, D. G. & Costerton, J. W. Biofilms as complex differentiated communities. Annu. Rev. Microbiol. 56, 187–209 (2002).

17. Motta, J.-P., Wallace, J. L., Buret, A. G., Deraison, C. & Vergnolle, N. Gastrointestinal biofilms in health and disease. Nat. Rev. Gastroenterol. Hepatol. 18, 314–334 (2021).

18. Buret, A. G. & Allain, T. Gut microbiota biofilms: From regulatory mechanisms to therapeutic targets. J. Exp. Med. 220, e20221743 (2023).

19. Smith, H. R. et al. Acidification-dependent suppression of C. difficile by enterococci in vitro. bioRxiv 2023.05.16.541032 (2023) doi:10.1101/2023.05.16.541032.

20. Permpoonpattana, P. et al. Functional characterization of Clostridium difficile spore coat proteins. J. Bacteriol. 195, 1492–1503 (2013).

21. Dubois, T. et al. A microbiota-generated bile salt induces biofilm formation in Clostridium difficile. npj Biofilms and Microbiomes 5, 14 (2019).

22. Flores-Mireles, A. L., Pinkner, J. S., Caparon, M. G. & Hultgren, S. J. EbpA vaccine antibodies block binding of Enterococcus faecalis to fibrinogen to prevent catheter-associated bladder infection in mice. Sci. Transl. Med. 6, 254ra127 (2014).

23. Schleifer, K. H. & Kilpper-Balz, R. Transfer of Streptococcus faecalis and Streptococcus faecium to the Genus Enterococcus nom. rev. as Enterococcus faecalis comb. nov. and Enterococcus faecium comb. nov. Int. J. Syst. Bacteriol. 34, 31–34 (1984).

24. Heydorn, A. et al. Quantification of biofilm structures by the novel computer program COMSTAT. Microbiology 146 (Pt 10), 2395–2407 (2000).

25. Davies, D. G. & Marques, C. N. H. A fatty acid messenger is responsible for inducing dispersion in microbial biofilms. J. Bacteriol. 191, 1393–1403 (2009).

26. Wetzel, D. & McBride, S. M. The Impact of pH on Clostridioides difficile Sporulation and Physiology. Appl. Environ. Microbiol. 86, (2020).

27. Sorg, J. A. & Sonenshein, A. L. Bile salts and glycine as cogerminants for Clostridium difficile spores. J. Bacteriol. 190, 2505–2512 (2008).

28. Arthur, M. & Quintiliani, R., Jr. Regulation of VanA- and VanB-type glycopeptide resistance in enterococci. Antimicrob. Agents Chemother. 45, 375–381 (2001).

29. Gardete, S. & Tomasz, A. Mechanisms of vancomycin resistance in Staphylococcus aureus. J. Clin. Invest. 124, 2836–2840 (2014).

30. Theriot, C. M. et al. Antibiotic-induced shifts in the mouse gut microbiome and metabolome increase susceptibility to Clostridium difficile infection. Nat. Commun. 5, 3114 (2014).

31. Rumbaugh, K. P. & Sauer, K. Biofilm dispersion. Nat. Rev. Microbiol. 18, 571–586 (2020).

32. Fleming, D. & Rumbaugh, K. The consequences of biofilm dispersal on the host. Sci. Rep. 8, 10738 (2018).

